# COPD lungs show an attached stratified mucus layer resembling the protective colonic mucus

**DOI:** 10.1101/205948

**Authors:** Joan Antoni Fernández-Blanco, Liisa Arike, Anna Ermund, Dalia Fakih, Ana M. Rodríguez-Piñeiro, Beatriz Martínez-Abad, Elin Skansebo, Sonya Jackson, James Root, Dave Singh, Christopher McCrae, Christopher M. Evans, Annika Åstrand, Gunnar C. Hansson

**Author notes:** Correspondence to: Gunnar C. Hansson, Department of Medical Biochemistry, Sahlgrenska Academy, University of Gothenburg, PO Box 440, S-405 30 Gothenburg, Sweden.

## Abstract

The respiratory tract is normally kept essentially free of bacteria by cilia-mediated mucus transport, but in chronic obstructive pulmonary disease (COPD) and cystic fibrosis (CF) mucus accumulates due to goblet cell hyperplasia and mucin overexpression. To address mechanisms behind the mucus accumulation, the elastase-induced mouse model was utilized. The proteomes of bronchoalveolar lavage fluid from elastase-induced mice and COPD patients showed similarities to each other and to colonic mucus. Lung mucus showed a striated, laminated appearance in the elastase-induced mice, COPD and CF, resembling that observed for colonic mucus. Less mucus obstruction was observed in mice lacking the Muc5b mucin. The accumulated mucus plugs of the elastase-induced mice were possible to wash out, but a mucus layer covering the epithelium remained attached to the surface goblet cells also after hypertonic saline washings as widely used in CF therapy. The results suggest that the lung can convert its mucus system into an attached mucus layer that protects the epithelium, similarly to the colon.

## Introduction

Stagnant mucus is a problem in several airway diseases including chronic obstructive pulmonary disease (COPD) (Smaldone *et al.*, 1993), cystic fibrosis (CF) (Regnis *et al.*, 1994), and asthma (Bateman *et al.*, 1983). COPD is a global health issue estimated to be the third leading cause of death worldwide in 2020 and the disease is an increasing economic and personal burden. It is characterized by restricted airflow and progressive loss of lung function and encompasses two major phenotypes, chronic bronchitis and emphysema, both caused by excessive inflammatory response to environmental noxious insults. Chronic bronchitis involves an increase in goblet cell number, enlarged submucosal glands, mucus hypersecretion, mucus plugging of the airways, and chronic cough (Vestbo *et al.*, 2013). Even COPD patients without chronic bronchitis have mucus obstruction of the small airways (Hogg *et al.*, 2004).

The mucociliary system keeps the respiratory tract low in bacteria and removes any inhaled particulate matter. Healthy mucus traps bacteria and particles that the mucus transports up to the larynx resulting in efficient cleaning. In numerous lung diseases, including COPD and CF, mucus stagnates and accumulates in the airways, leading to bacterial colonization, airway inflammation tissue destruction, and ultimately respiratory failure. Mechanisms of mucociliary dysfunction include the aberrant overproduction of mucus components leading to accumulation on the airway surfaces. Mucus is made up by many different components, but mucins are the major macromolecular constituents. In human and pig airways, there are two major gel-forming mucins, the MUC5B from the glands and the MUC5AC from the surface goblet cells. The glands form MUC5B mucin bundles that under normal conditions sweep the tracheobronchial surface to remove debris and bacteria (Ermund *et al.*, 2017). In rodents, which lack submucosal glands, both Muc5b and Muc5ac mucins are made by the surface goblet cells. Also in rodents, the Muc5b mucin is essential for keeping the mouse lungs clean under normal conditions (Roy *et al.*, 2014).

There are several rodent models of chronic airway disease. For example, models induced by ovalbumin (Kung *et al.*, 1994), IL–13 (Kuperman *et al.*, 2002), LPS instillation, or cigarette smoke (Shapiro, 2000) are used. Airway instillation of various forms of elastase, for example neutrophil elastase or porcine pancreatic elastase (PPE), in mice, rats, guinea pigs, dogs or ferrets has been widely used as models of COPD and emphysema (Conlon *et al.*, 2016;Gross *et al.*, 1964). This can also be extended to CF since neutrophils accumulate in CF lungs (Birrer *et al.*, 1994). Based on the importance of elastase in chronic lung diseases and the fast onset of goblet cell hyperplasia we focused on the elastase model. PPE is instilled intranasally twice with a 7 day interval and animals analyzed two weeks after the first instillation. We have analyzed this model for goblet cell hyperplasia, mucus accumulation, inflammation, and mucus proteins and compared the mucus proteome to human airway disease and colon mucus. The importance of the Muc5ac and Muc5b mucins were addressed by using knockout mice. We tested a common treatment for mucus accumulation in the airways, hypertonic saline, and its ability to detach the accumulated mucus. Collectively, our results show a striking similarity between the accumulated mucus in the respiratory tract of mice and COPD patients with that of the gastrointestinal tract.

## Results

### PPE treatment causes goblet cell hyperplasia and airway plugging

Mice intranasally exposed to saline largely lacked Alcian blue/Periodic acid-Schiff (AB/PAS) positive cells in the airways whereas a marked increase in AB/PAS positive cells within the bronchi and bronchioles was observed in PPE-exposed mice and not after saline-or PPE-inactivated treatment (Fig 1A-D, S1). Whole mouse lungs stained with AB/PAS showed that the hyperplasia had a patchy distribution, affecting bronchi and proximal bronchioles with both goblet cell hyperplasia and mucus plugging (Fig 1CD). This was further illustrated by transmission electron microscopy (Fig S2A). Histopathology evaluation of inflated lungs from PPE-challenged mice revealed a mild perivascular and peribronchiolar lymphocytic inflammation relative to control mice (Fig 1E, S2B). These alterations were accompanied by a patchy destruction of the inter-alveolar septa with moderate to severe multi-focal loss of tissue connectivity as in emphysema. Mucus obstruction was quantified in AB/PAS-stained sections, presented as percent airway obstruction, showing a clear increased airway obstruction in PPE-exposed mice Fig 1F where the smallest airways were more prone to develop severe mucus obstruction (Fig S3). Bronchoalveolar lavage fluid (BALF) from PPE-exposed mice showed higher immune cell counts with a 16-fold increase in the total number of white blood cells, consisting of increased amounts of neutrophils, eosinophils, macrophages/monocytes and lymphocytes (Fig 1G). The increased number of inflammatory cells was accompanied by increased levels of cytokines, chemo attractants, chemokines, and EGF (Fig S2). In summary, PPE treatment caused increase in goblet cell numbers, mucus accumulation and immune cell infiltration as in COPD.

**Figure 1.**
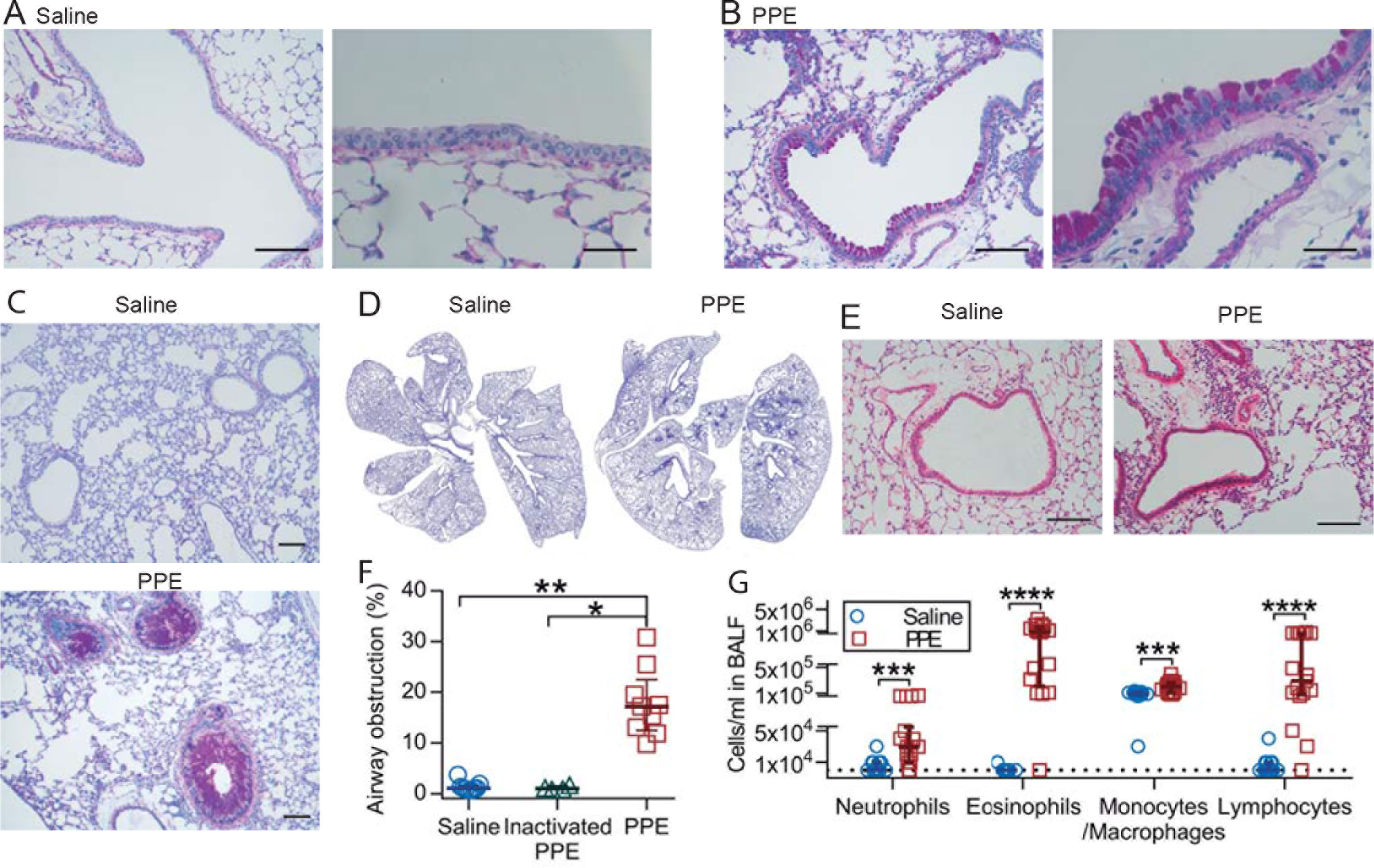
Mice exposed to PPE show goblet cell hyperplasia/metaplasia and mucus accumulation in the airways. **A** Alcian blue/Periodic acid-Schiff (AB/PAS) stained tissue section of airways mice exposed intranasally to saline (vehicle). **B** Alcian blue/Periodic acid–Schiff (AB/PAS) stained tissue section of airways mice exposed intranasally to PPE showing a marked increase in AB/PAS positive cells (A and B) Representative images of 4–5 animals/group. Scale bar, 100 μm (left panels) or 33 μm (right panels). **CD** High and low high magnification images of lungs from mice exposed to saline or PPE. Scale bar in C, 100 μm. **E** Paraffin sections stained with H&Ε revealed few immune cells around intrapulmonary airways and intact alveoli from saline–instilled mice. Mild perivascular and peribronchiolar lymphocytic infiltration and damaged alveoli was detected in PPE-challenged mice. **F** Airway obstruction presented by measuring the percentage of airway luminal area containing AB/PAS–stained material in one entire lung section per animal, n = 4–9 animals/group, median ± interquartile range, saline vs. PPE: P = 0.001**, inactivatedPPE vs. PPE: P = 0.01*, Kruskal-Wallis with Dunn´s post hoc test. **G** Differential white blood cell counts in BALF of vehicle– and PPE-exposed mice, n = 9–17 animals/group, data presented as median ± interquartile range, neutrophils P =0.0008***, eosinophils P < 0.0001****, macroaphages P = 0.0005***and lymphocytes P 0.0001****, Mann-Whitney U test.

### Proteome of mouse bronchoalveolar lavage fluid and mucus plugs

Total BALF samples collected from saline-or PPE-induced mice were analyzed by mass spectrometry based proteomics. To remove cells from BALF, samples were centrifuged and the supernatants analyzed where we observed 349 proteins in controls and 408 in PPE-exposed animals (Table 1, S1). Some of the major proteins including the club cell secreted protein uteroglobin (Scgb1a1) serotransferrin (Tf), and α-1-antitrypsin (Serpin1ad) were common for saline or PPE mice, whereas a relative increase in mucus-related proteins including Clca1, Chitinase-like proteins Chi3l3 and Chi3l4 were observed in the PPE mice (Fig 2AB). Absolute quantification of the gel-forming mucins Muc5b and Muc5ac corroborated the observed PPE-mediated goblet cell hyperplasia where Muc5ac, not detected in controls, reached concentrations similar to that of Muc5b (Fig 2C, Table S2). Muc5b levels were increased 24-fold compared to basal conditions. To better understand the proteins accumulated by PPE, we isolated mucus plugs from BALF after staining with Alcian blue (Fig 2D). The major components of the airway plugs were similar to those of the BALF, but had an even higher relative level of Clca1 (Fig 2E,Table 1, S3). In addition, the plugs also contained high proportions of the Bpifb1 (also known as Lplunc1), Muc5ac, Muc5b and Fcgbp. Absolute quantification of Muc5ac and Muc5b in the plugs revealed significantly higher levels of Muc5b relative Muc5ac, in contrast to the BALF were more similar levels were found in PPE-induced mice (Fig 2).

**Figure 2.**
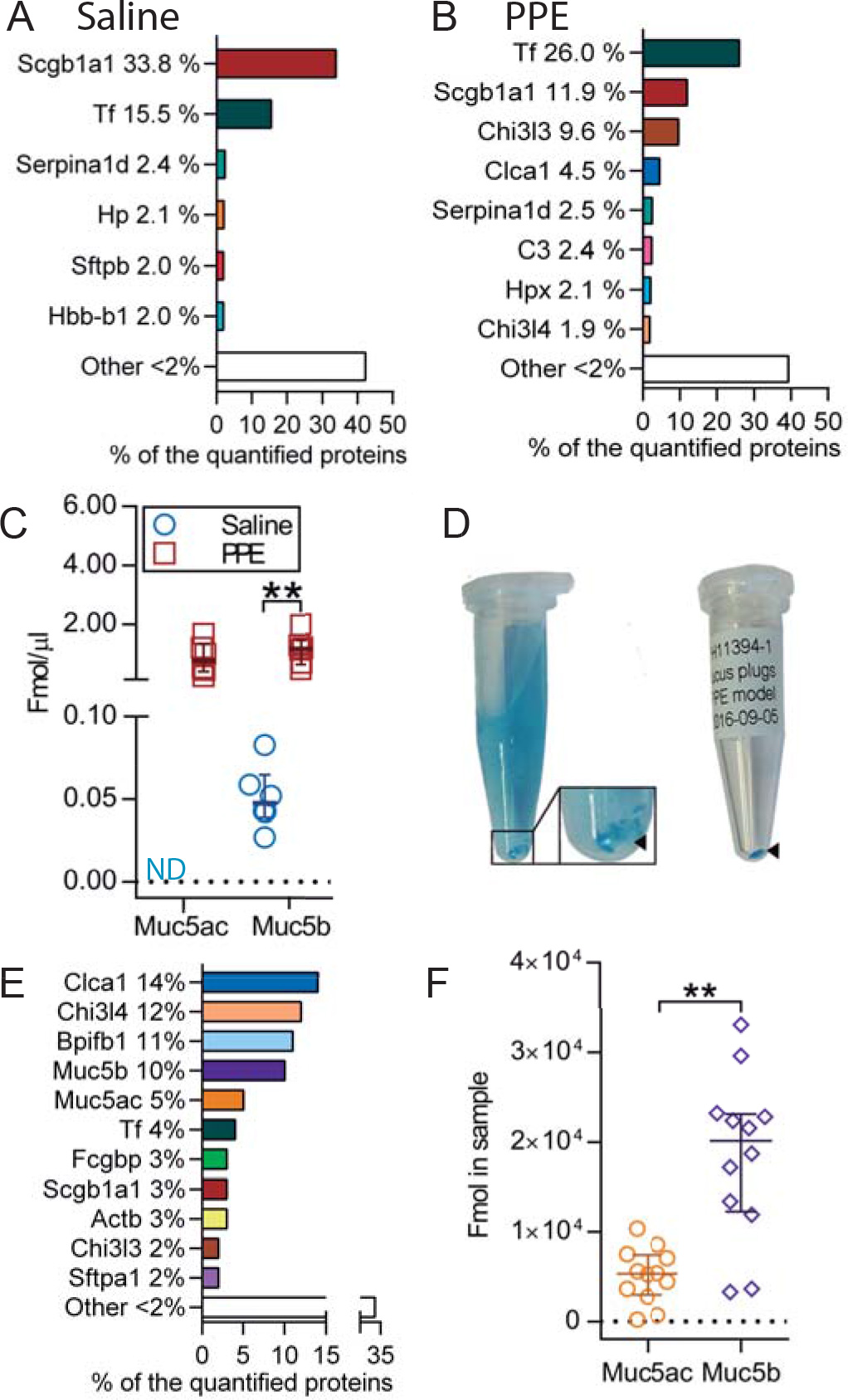
PPE administration alters BALF proteome. **A**, **B** Major proteins in BALF supernatants obtained from lungs lavaged with PBS in animals exposed to saline (A) and PPE (B) analyzed by mass spectrometry proteomics. **C** Absolute amounts of Muc5ac and Muc5b in BALF supernatant from saline– and PPE-exposed mice analyzed by mass spectrometry. ND means not detected, n = 6 animals/group in A–C, data presented as median ± interquartile range, P = 0.002 **, Mann-Whitney U test. **D** Isolation of mucus plugs from BALF in a PPE–treated mouse after lavage with an Alcian blue solution. Blue–stained mucus plugs (arrowhead) were obtained and transferred to a dry tube (right). **E** The most abundant proteins in mucus plugs detected by proteomics. **F** Absolute quantification sof Muc5ac and Muc5b protein amounts in mucus plugs by mass spectrometry, data presented as median ± interquartile range, n = 12 animals in E and F, P = 0.0011**, Mann–Whitney U test. G Label–free quantification (LFQ) of the most changed proteins in isolated airway epithelial cell after saline or PPE treatment, n = 7pools/group with 3 mice/pool, data presented as median ± interquartile range, P= 0.017*, 0.006**, 0.0006***, Mann-Whitney U test. Scgb1a1: uteroglobin; Tf: serotransferrin; Serpina1d: alpha–1–antitrypsin; Hp: haptoglobin; Sftpb: surfactant protein B; Hbb–0b1: hemoglobin subunit beta–1; Chi3l1/3/4: chitinase–1/3/4–like protein; Clca1; C3: complement C3; Hpx: hemopexin; Chi3l4: chitinase–3–like protein 4; Bpifb1: BPI fold-containing family B member 1; Fcgbp; Actb: actin cytoplasmic; Sftpa1: pulmonary surfactant–associated protein A.

**Table 1.**
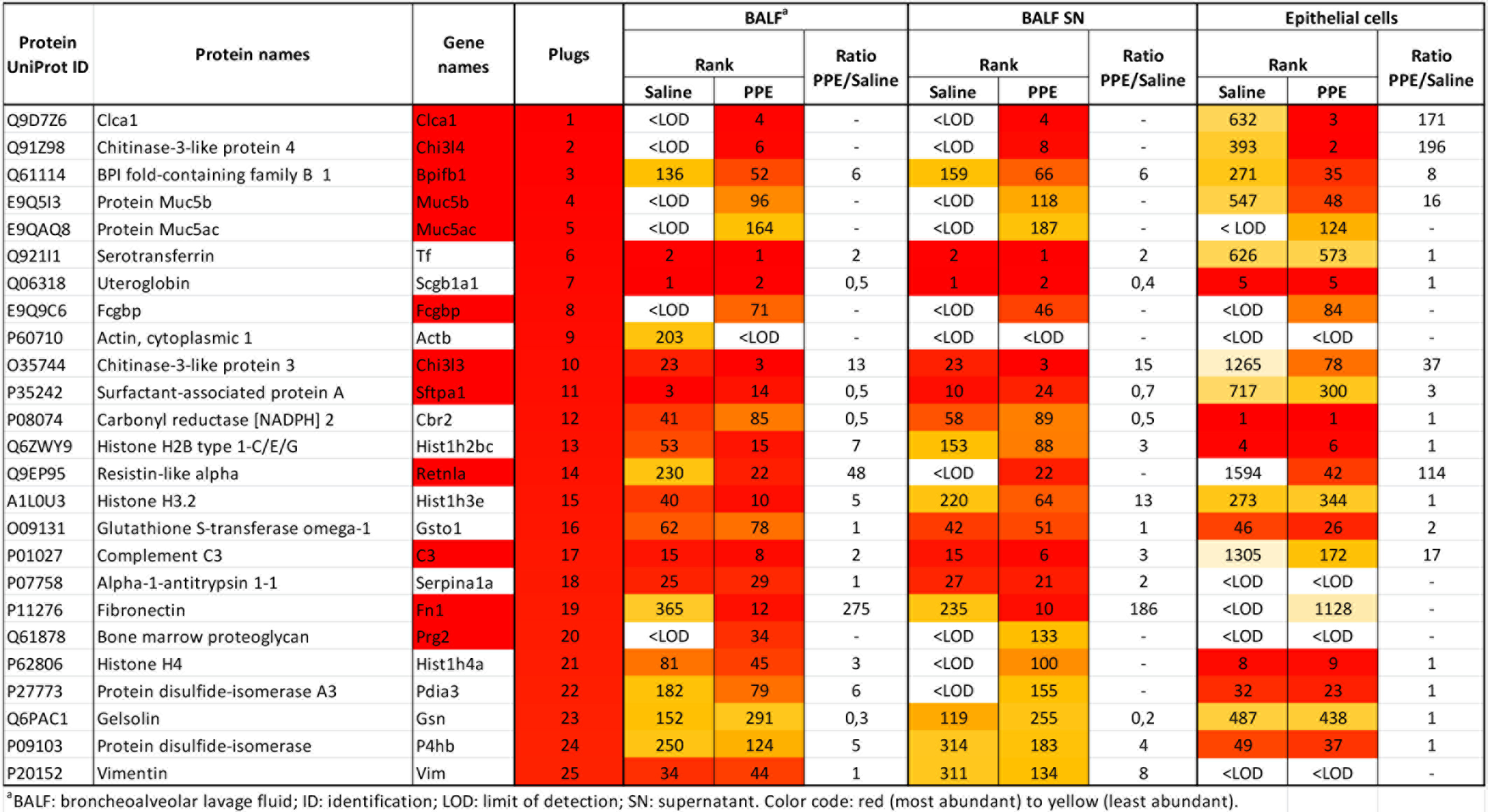
Most abundant proteins (excluding albumin) detected by mass-spectrometry in airway mucus plugs obtained from mice exposed to PPE and changes induced in the epithelial cell
and BALF proteome in vivo.

### Proteome of isolated mouse airway epithelial cells

To further illustrate the PPE effect, we isolated airway epithelial cells and analyzed their proteome (Table 1,Table S4). The isolation procedure was effective as shown by mouse lung sections stained with H&E before and after epithelial removal(Fig 3A). Heat map clustering of the proteomic results showed that the epithelial cells of the PPE-exposed mice were clearly separated from the controls with the most noteworthy differences around the Muc5b and Muc5ac nodes (Fig 3B). Chi3l4 and Clca1 were among the most abundant proteins in PPE–exposed mice (Fig 3CD. These and the other mucus-related proteins found in PPE-induced BALF and mucus plugs were also increased in epithelial cells, but the chitinase-like proteins were relatively more increased in the PPE-induced epithelial cells (Fig 3E). Furthermore, the increased levels of immune response proteins in observed in BALF were also found in the epithelial cell fraction (Table S1,S3). Absolute quantification revealed, in agreement with the BALF, that Muc5ac was only detected after PPE treatment and that Muc5b levels were increased, but in this case only 7.6 times (Fig 3F). Muc5b was always more abundant than Muc5ac, in mucus plugs with ratio Muc5ac/Muc5b of 0.3, in BALF supernatant (BALF SN) of 0.7, and in epithelial cells 0.5 (Fig 3G).

To support the proteomics and to analyze the tissue localization of key proteins immunostaining was performed (Fig. S4, S5). After PPE instillation, Muc5b, Muc5ac, Clca1, and Fcgbp were all found to be increased in the epithelial cells of both bronchi and bronchioles and all these proteins were accumulated in mucus plugs obstructing the airways. The few Muc5ac-positive epithelial cells detected at basal conditions also expressed Muc5b and after PPE a higher proportion of cells co-expressing Muc5ac and Muc5b (Fig S4E).

**Figure 3.**
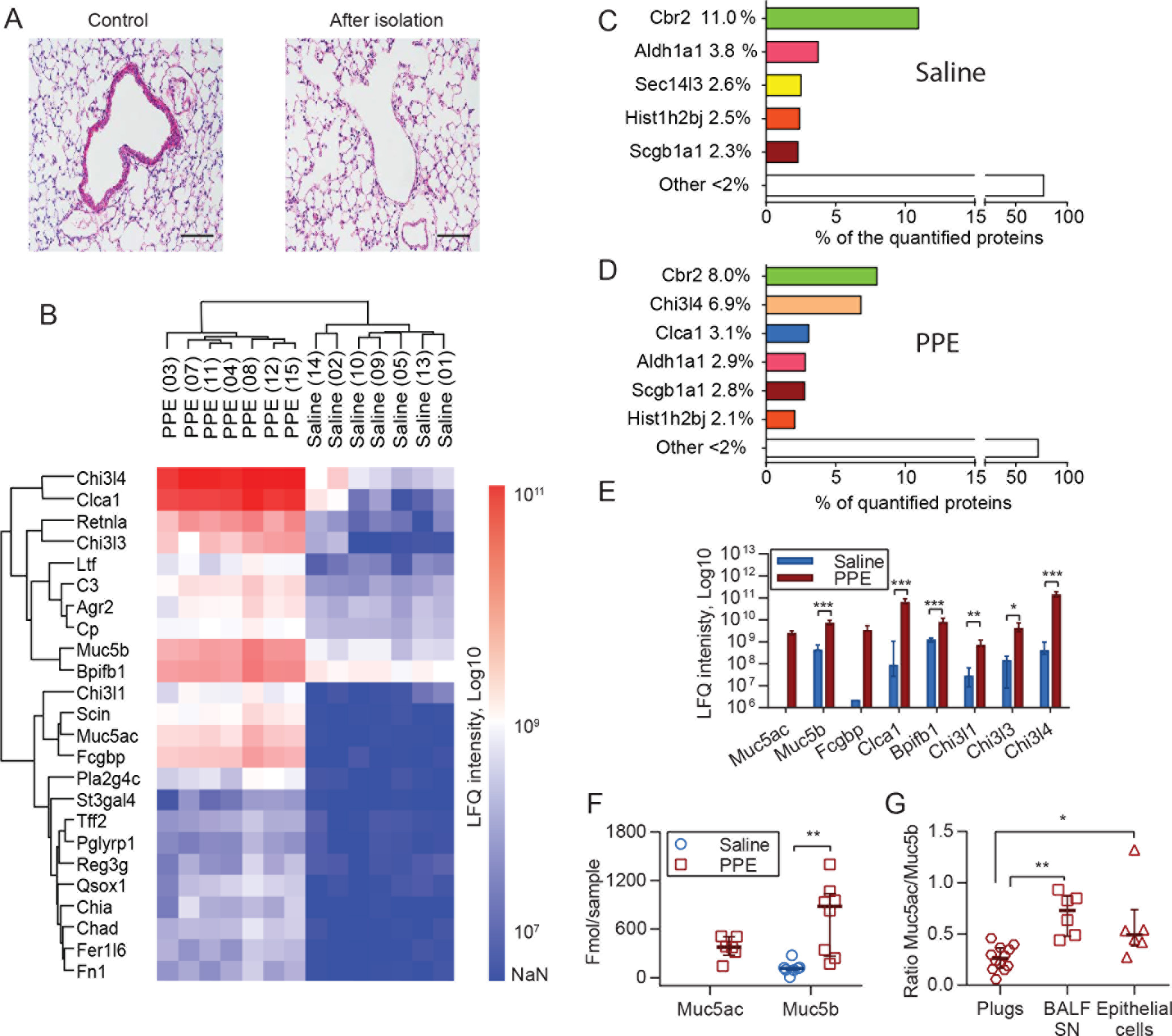
Airway epithelial cell proteome changes after PPE administration. **A** H&E stained section of bronchi before (left) and after epithelial cell isolation (right). **B** Heat map clustering of most changed proteins in airway epithelial cell proteome after PPE treatment. **C**, **D** The most abundant proteins in airway epithelial cells detected by label-free proteomics after saline (C) and PPE (D) instillation. **E** Label–free quantification (LFQ) of the most changed proteins after PPE treatment, n =7 pools/group with 3 mice/pool, data presented as median ± interquartile range, P=0.017*, 0.006**, 0.0006***, Mann-Whitney U test. **F** Absolute quantification of Muc5ac and Muc5b with isotopically labelled peptides by mass spectrometry, n = 8 pools/group with 3 mice/pool, data presented as median ± interquartile range, P = 0.001**, Mann–Whitney U test. **G** Ratio of Muc5ac to Muc5b in isolated mucus plugs, BALF supernatants (BALF SN) and epithelial cells, n = 6–12 samples/group. Data presented as median ± interquartile range, P = 0.04*, 0.0016**, Kruskal–Wallis and Dunns post hoc test. Chi3l4: chitinase–3– like protein 4; Clca1: chloride channel accessory 1; Retnla: Resistin-like alpha; Chi3l3:chitinase–3–like protein 3; Ltf: Lactotransferrin; C3: complement C3; Agr2: Anterior gradient protein 2 homolog; Cp: Ceruloplasmin; Bpifb1: BPI fold–containing family B member 1; Chi3l1: Chitinase-3-like protein 1; Scin: Adseverin; Fcgbp; Pla2g4c: Pla2g4cprotein; St3gal4: Beta–galactoside alpha-2,3-sialyltransferase 4; Tff2: Trefoil factor 2;Pglyrp1: Peptidoglycan recognition protein 1; Reg3g: Regenerating islet-derived protein3–gamma; Qsox1: Sulfhydryl oxidase 1; Chia: Acidic mammalian chitinase; Chad:Chondroadherin; Fer1l6: Fer–1–like 6; Fn1: Fibronectin; Cbr2: Carbonyl reductase [NADPH] 2; Aldh1a1: Retinal dehydrogenase 1; Sec14l3: SEC14-like 3; Hist1h2bj: Histone H2B; Scgb1a1: uteroglobin.

### Human bronchoalveolar fluid proteome resembles that of the mouse model

To compare the PPE-induced mouse model with the human disease COPD, we studied human BALF by mass spectrometry based proteomics from never smokers, asymptomatic smokers and COPD patients (Table S5). Proteomics analysis revealed fewer proteins in the never smokers group as compared to asymptomatic smokers and COPD patients (Fig 4A, Table S6). Principal component analysis showed a tight cluster for the never smokers and a slightly larger for the asymptomatic smokers (Fig 4B). Interestingly, the COPD patients were distributed over the entire plot suggesting a relatively large variation between the individuals. Absolute quantification of a subset of mucus-related proteins revealed increased amounts of AGR2, DMBT1, FCGBP, PSCA, TFF3, TGM2 and ZG16B in COPD patients and a tendency towards increased levels of these proteins in asymptomatic smokers (Fig 4C, S6, S7 and Table S7). In accordance with the PPE model, MUC5AC and MUC5B were increased in COPD patients and the ratio MUC5AC to MUC5B changed from 0.3 in never smokers to closer to 1.5 in asymptomatic smokers and 0.5 in COPD patients (Fig 4D). In contrast to the mouse model where Clca1 was one of the most abundant proteins after PPE treatment, CLCA1 was not detected in human BALF. In summary, the human COPD BALF showed large similarities with the PPE mice, although there are some species specific differences.

**Figure 4.**
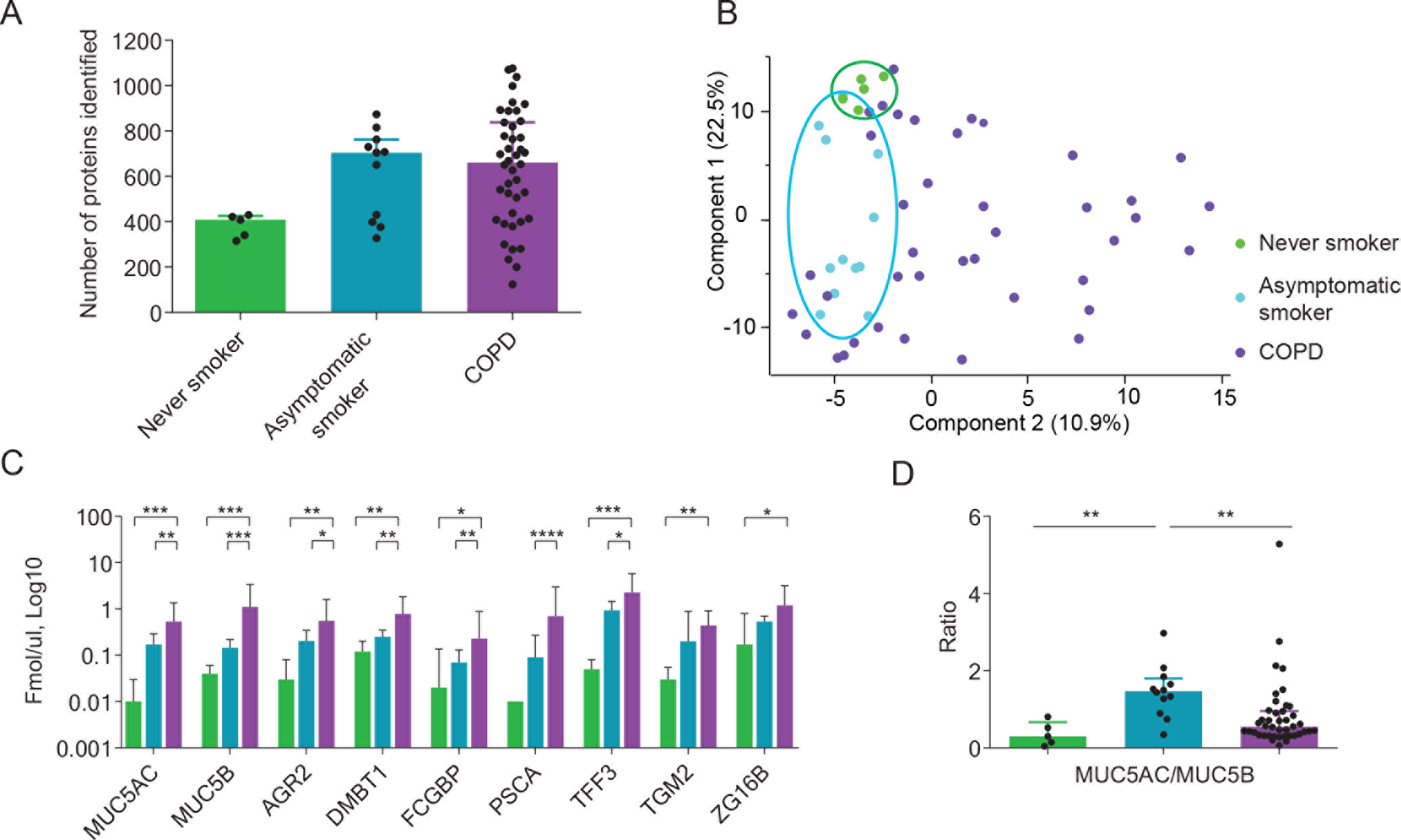
Altered proteome in BALF from never smokers (n=5), asymptomatic smokers (n=12) and chronic obstructive pulmonary disease (COPD) patients (n=42). **A** Number of proteins identified in BALF in each sample group, n = 5–42, data presented as medians interquartile ranges. **B** Principal component analysis. **C** Absolute quantification with isotopically labelled peptides by mass spectrometry, n = 5–42, data presented as medians interquartile ranges, P = <0.05*, <0.01**, 0.001***,0.0001****/0.0001, Kruskal-Wallis and Dunns post hoc test. **D** Ratio of MUC5AC to MUC5B calculated from absolute quantification data in (C), n =5-42 patients/group, data presented as medians interquartile ranges, P=0.01**/0.01, Kruskal-Wallis and Dunns post hoc test. AGR2: Anterior gradient protein 2 homolog;DMBT1: Deleted in malignant brain tumors 1; FCGBP: Fc fragment of IgG binding protein; PSCA: Prostate stem cell antigen; TFF3: Trefoil factor 3; TGM2:Transglutaminase-2; ZG16B: Zymogen granule protein 16 homolog B.

### Airway mucus in elastase-exposed mice, COPD, and CF appeared stratified as in colon

Tissue fixation using dry Carnoy has been shown to preserve the mucus structure of the intestine (Johansson *et al.*, 2008). When the PPE-exposed mouse airways were fixed this way and stained by AB/PAS, the mucus in had a stratified, lamellar appearance (Fig 5A). When mucus was stained with antibodies against Muc5b (green) and Muc5ac (red), the two mucins appeared to form partly separate layers covering the epithelium (Fig 5B). A stratified mucus appearance was not been observed in control lungs, but resembled the inner mucus layer of the normal mouse distal colon (Fig 5C). Similar stratified mucus was also observed in lungs from human COPD (Fig 5D) and CF (Fig 5D) patients that had undergone lung transplantation. Thus, the mucus accumulated in the PPE model resembled the two chronic lung diseases with an appearance of the mucus similar to the stratified inner colon mucus.

**Figure 5.**
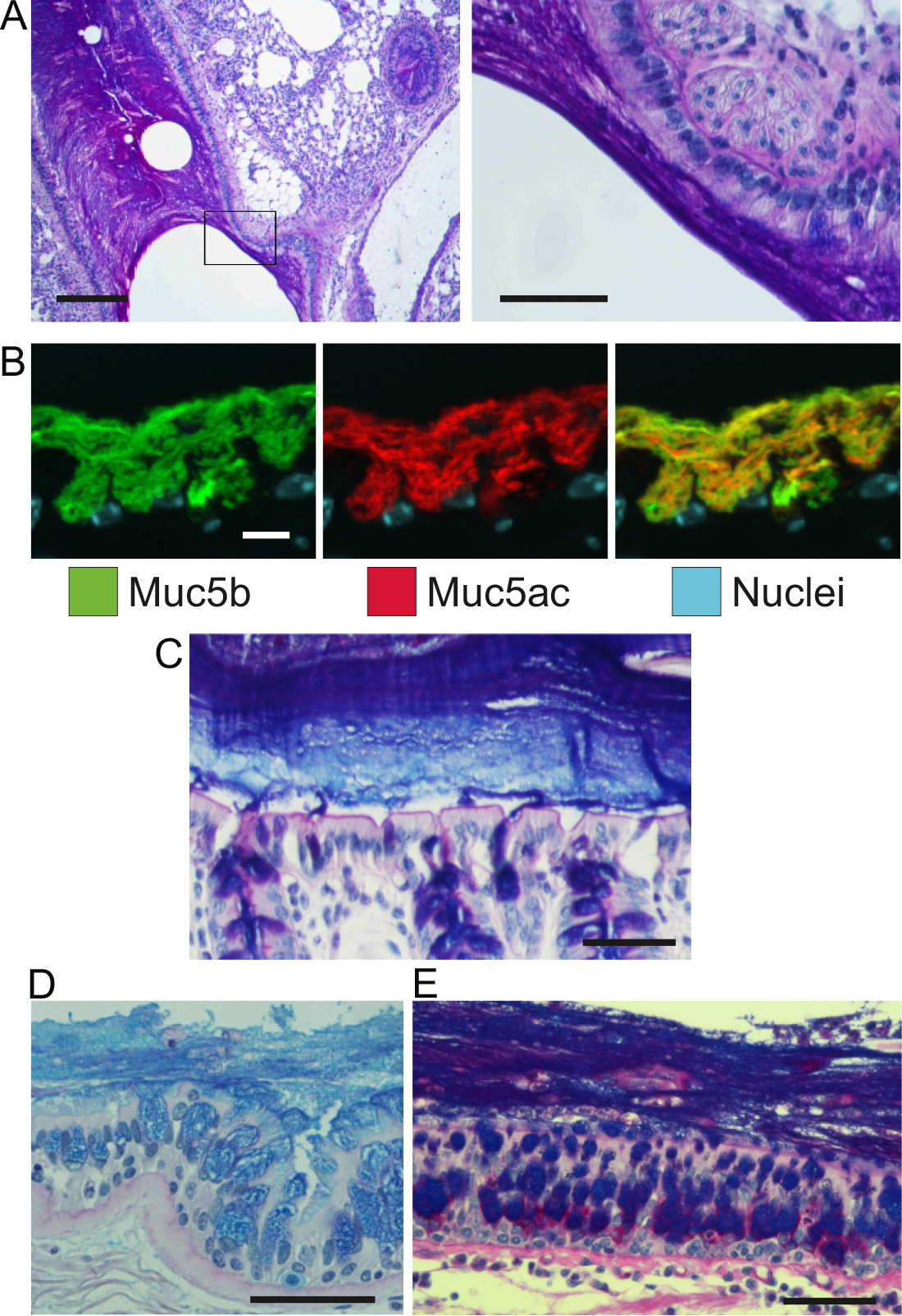
Airway mucus accumulated after PPE administration in mice has a stratified appearance. **A** Lower (left) and higher magnification (right) images of an AB/PAS stained paraffin section showing bronchial walls lined by mucus with lamellar appearance in a mouse exposed to PPE. Representative of 9 animals, scale bars: left 200 μm, right 50 μm. **B** Paraffin section from a mouse exposed to PPE with specific antibodies against Muc5b (green) and Muc5ac (red) in addition to nuclear stain (blue) shows that this stratified structure consisted of interchanging layers of Muc5b and Muc5ac, without actual mixing of the two mucins. Representative of 3 animals, scale bar 10 μm. **C** The stratified appearance of mucus has been observed in the distal colon where the mucus is organized to protect the epithelium from bacterial contact in humans as well as mice. Representative image showing mouse distal colon in an AB/PAS-stained paraffin section, scale bar 50 μm. In mice as well as humans, mucus does not accumulate in the airways in control subjects. **D**, **E** However, in airway paraffin sections from patients with COPD (D) and cystic fibrosis (E) stained with AB/PAS, stratified mucus can be observed covering the epithelium. Scale bars in D and E 50 μm.

### Mucus is anchored to the goblet cells in elastase-exposed mouse airways and cannot be completely washed away

To analyze if the accumulated mucus was attached to the epithelium in PPE-instilled mouse airways as in the CF mouse ileum (15), we lavaged PPE-exposed lungs twice with PBS. The mucus seemed less dense after this treatment as assessed from Carnoy-fixed tissue sections stained with AB/PAS (Fig 6A). Using these sections, we analyzed the percentage of total airways that were obstructed by mucus in non-washed and in PBS-lavaged lungs. There was a tendency for less airway obstruction in the PPE-exposed mouse airways after PBS lavage (Fig 6B). We also estimated the proportion of the airway surface that was covered by mucus without and after washing. As there were essentially no mucus covering the epithelium after saline treatment, there was substantially more epithelial surface covered by mucus in PPE-exposed airways (Fig 6C). Interestingly, there was no tendency of the PBS lavage to remove any of the mucus on the surface of PPE-exposed airways.

As the accumulated mucus contained both the Muc5b and Muc5ac mucins, we next addressed the individual importance of these mucins using knockout animals. The Muc5b^-/-^ mice showed significant reduced mucus obstruction whereas the Muc5ac^-/-^ only showed a tendency of reduced mucus obstruction (Fig 6D).

We have previously shown that hypertonic saline treatment can detach the abnormally attached mucus of the CF mouse ileum (11). This treatment is also approved for use in CF patients. To evaluate if hypertonic saline could remove the attached mucus in the PPE-exposed lungs, we instilled 7% hypertonic saline twice into the lungs, left it for 20 min each time, and aspirated the remaining liquid. As a control, we instilled PBS twice for 20 min. The hypertonic saline treatment caused a significant decrease in the percentage of airways that were obstructed by mucus (Fig 6E). When the percentage of airway surface covered by mucus was evaluated, there was almost no reduced epithelial surface covered by mucus (Fig 6F). This suggests that the 7% hypertonic saline can remove most of the mucus plugging, but that the surface attached mucus still remained.

To further illustrate how the mucus was attached, we stained Carnoy fixed tissue sections taken after two times 20 min PBS washings with antibodies against Muc5ac and Muc5b and analyzed this by confocal microscopy (Fig 6G). The epithelial cell surface was covered by a mucus layer composed of a mixture of the Muc5ac and Muc5b mucins. Both mucins seemed to be retained and anchored in every goblet cell. As it has been proposed that the stagnated lung mucus is attached by entangling into the cilia, we also stained tissue sections for cilia using an anti-tubulin antibody (Fig 6H). The image shows how the cilia extended in between the Muc5b stained mucus and do not suggest that the cilia were entangled with the mucus. Together these results suggest that the accumulated mucus in the PPE mouse airways is attached to the goblet cells, just as has been observed in the CF mouse intestine (Gustafsson *et al.*, 2012).

**Figure 6.**
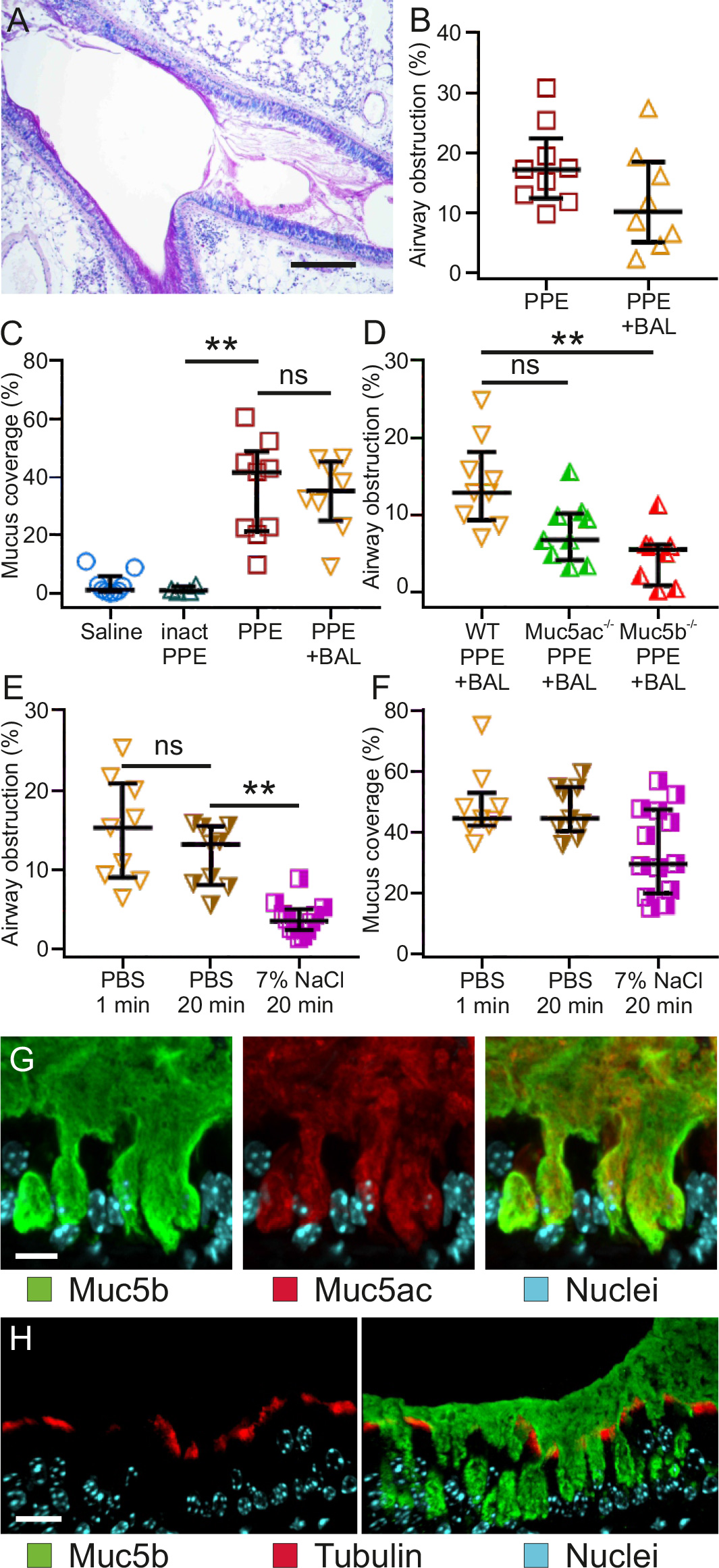
Accumulated airway PPE mucus is attached to the epithelium and can be removed by hypertonic saline. **A** Representative image of an airway from a mouse exposed to PPE and lavaged with PBS (BAL; 2 times, 1 min). Paraffin section stained with AB/PAS. Representative of 8 animals. Scale bar 200 μm. Note the lighter-seeming AB/PAS staining and the mucus still covering the epithelial surface. **B** Airway obstruction in percent measured as airway luminal area containing AB/PASstained material in an entire lung section per animal. n = 8−9 animals/group. 695−729 airway sections/group. BAL was performed by washing 2 times 1 min with PBS. Results presented as median ± interquartile range. **C** Graph showing the large increase in percentage of airway surface covered byAB/PAS/stained material in mice exposed to PPE compared to inactivated PPE. The inactivated PPE gives no increase in surface area covered by mucus, corroborating that the enzymatic activity of PPE induces the mucus accumulation. In addition, two 1 min washes with PBS did not affect the percentage of epithelium covered by mucus after PPE treatment, n = 4–9 animals/group, median ± interquartile range, ** p = 0.009, Kruskal-Wallis and Dunnfs multiple comparisons test. **D** Airway obstruction after BAL (2x1 minute) in PPE treated Muc5ac^-/-^ and Muc5b^-/-^ as compared to WT, n=8−9, **p= 0.011.**E** Airway obstruction was reduced after two times 20 min treatment with7% saline compared to two times 20 min wash with PBS measured in AB/PAS stained paraffin sections, n = 9−12 animals, ** p = 0.002, Kruskal-Wallis and Dunn´s multiple comparisons test. **F** The percent of the epithelium covered by mucus was not affected by two times 20 min 7% saline treatment measured in AB/PAS stained paraffin sections, n = 9−13 animals, Kruskal-Wallis and DunnLs multiple comparisons test. **G** Immunostaining of paraffin section as in A but with specific antibodies against Muc5b (green) and Muc5ac (red) in addition to nuclear stain (blue). Both Muc5b and Muc5ac seem to be attached inside goblet cells. Representative of 4 mice/group, scale 20 μm. **H** Immunostaining of a paraffin section from a PPE exposed mouse with specific antibodies against Muc5b (green) and tubulin (red) to visualize cilia in addition to nuclear stain (blue). The cilia are not compressed by the accumulated mucus. Representative of 3 mice, scale 20 μm.

### Discussion

Airway diseases such as COPD and CF involve impaired mucociliary clearance which leads to mucus accumulation, bacterial retention, bacterial colonization, tissue destruction,and respiratory failure. The mechanisms behind mucus retention and plugging of the airways are despite considerable efforts largely unknown. The use of human bronchial epithelial (HBE) cultures has provided important information on the mucus properties in CF, but the relevance of this model for lung disease is not fully understood (Matsui *et al.*,1998). With the aim to utilize a convenient animal model for goblet cell hyperplasia and mucus retention, we chose intranasal instillation of elastase in mice. An increased amount of neutrophils and their release of elastase is considered a hallmark for the development of COPD (Hoenderdos and Condliffe, 2013). We instilled porcine pancreatic elastase (PPE) twice with an interval of 7 days before analyzing the animals after 7 additional days. We studied the elastase-treated mice by histological analyses, immune cells, inflammatory mediators, and especially proteomics of BALF and epithelial cells. The PPE exposure activated pathways resulting in a dramatic goblet cell hyperplasia and metaplasia. The mouse lungs showed mucus accumulation with a patchy profile characteristic for human chronic bronchitis and COPD. Severe mucus obstruction was observed especially in airways with smaller diameter corresponding to peripheral bronchi and proximal bronchioles as is commonly observed in human lung disease.

Increased inflammation is characteristic for the airway diseases chronic bronchitis, COPD, CF and asthma. The inflammatory response in PPE-exposed mice was dominated by eosinophils, but neutrophils, macrophages/monocytes and lymphocytes were also increased. The mouse lungs are different from the humans as they essentially lack submucosal glands and thus the PPE mouse model cannot be a model for any particular human disease, but has traits of chronic bronchitis, COPD, CF, and possibly also late and severe asthma. Among the chemokines upregulated in PPE-exposed mice was KC, a murine homologue of IL–8. Higher concentrations of IL-8 have been detected in sputum from COPD patients than asthma patients and IL-8 has been correlated with poorer lung function measured as FEV_1_ in COPD patients (Yamamoto *et al.*, 1997). In asthma, but also in some patients with COPD, the lung mucosa is infiltrated by eosinophils and Th2 cells contribute to tissue damage. The Th2 cells produce cytokines, such as IL–4,IL–5 and IL–13 (Lukacs, 2001), of which IL–4 and IL–5 were also increased in the PPE model. Thus, the PPE model showed immunological characteristics that are relevant for studying mucus accumulation in airway disease.

We have previously performed extensive mass spectrometric proteomic studies of the mouse gastrointestinal mucus proteome (Johansson *et al.*, 2009;Rodriguez-Pineiro *et al.*,2013). In normal mouse colon mucus, Clca1, Fcgbp, Agr2, and Zg16 are among the most abundant proteins and similar in amount to the main intestinal gel-forming mucin, Muc2. All these proteins are made by the colonic goblet cells. The proteome of BALF, mucus plugs as well as epithelial cells from the lungs of PPE-exposed mice showed increased levels of all these proteins as compared to controls except for Zg16 that is not present in mouse lungs. Normal mouse lungs rarely express the gel-forming Muc5ac mucin, but this was dramatically induced together with Muc5b after PPE-treatment. Also other proteins, like Bpifb1, Chi3l1, Chi3l3, and Chi3l4 were highly increased. Bpifb1, also called SPLUNC1, has been shown to be increased in sputum of COPD as well as in CF patients compared to non-smokers (Bingle *et al.*, 2012;Gao *et al.*, 2015).

Studies on human BALF samples from COPD patients by proteomic were used to evaluate the similarities with that of the PPE-exposed mouse. Interestingly, principal component analysis of the mass spectrometry results clearly distinguished never smokers, asymptomatic smokers and COPD patients. The never smokers and the asymptomatic smokers clustered well together, whilst the COPD patients spread over the whole graph suggesting a large heterogeneity. Comparing the absolute amounts of selected proteins, known to be associated with mucus, showed that all were increased in COPD as compared to never smokers with the asymptomatic smokers in between. In accordance with the PPE-exposed mice, the two mucins (MUC5AC and MUC5B), AGR2, and FCGBP were all increased in COPD. Another known goblet cell product, trefoil factor 3 (TFF3) was found in both the human intestine and airways (Podolsky *et al.*, 1993;Wiede *et al.*, 1999). In the airways DMBT1, a scaffolding protein involved in binding bacteria facilitating their removal, was also increased in COPD BALF (Madsen *et al.*, 2010). The prostate stem cell antigen (PSCA) was increased in COPD patients compared to asymptomatic and never smokers. Interestingly, PSCA has been suggested to act as a negative regulator of the α7 nicotinic acetylcholine receptors and by this inhibiting nicotinic signaling (Fu *et al.*, 2015), something that might be related to nicotine intake as all COPD patients were smokers. Increased amounts of transglutaminase 2 (TGM2) has previously been observed in COPD (Ohlmeier *et al.*, 2016), indicating that also lung mucins can be covalently crosslinked as observed in colon (Recktenwald and Hansson, 2016).

Comparing the lung BALF proteome between humans and mice revealed some discrepancies probably related to species differences. The chitinase-like proteins were more abundant in mice. Still, the chitinase 3-like protein (CHI3L1) found in airway epithelial cells is a marker of human asthma severity and declined lung function (Komi *et al.*, 2016). The Clca1 protein, a typical colon mucus protein, was dramatically increased in the PPE-exposed mice, but not in human BALF. Clca1 was first suggested to be related to chloride ion channels, but more recent studies have shown that it is a protease that might be involved in mucus turnover (Lee *et al.*, 2015). The ZG16 protein produced by goblet cells is abundant in mouse and human colonic mucus where it protects the epithelial cells by aggregating gram-positive bacteria and by this translocating them further away from the epithelial cell surface (Bergström *et al.*, 2016). This protein is absent in the human BALF, but a similar protein named ZG16B, was highly increased in COPD patients. However, this protein was not present in mice.

Increased levels of the two gel-forming mucins MUC5AC and MUC5B are probably the most typical alteration in the human lung diseases chronic bronchitis, COPD, CF and asthma as well as in all animal models (Thornton *et al.*, 2008). This was also common for both the PPE mouse model and human COPD samples studied herein. This common feature is irrespective of the different organization of the normal mucus producing cells in mice and humans where the latter have submucosal glands that make long MUC5B bundles (Ermund *et al.*, 2017). In normal mice the surface goblet cells more or less only produce the Muc5b mucin, but there is a dramatic increase in the Muc5ac produced upon the induced goblet cell hyperplasia. The MUC5B mucin forms linear polymers responsible for normal clearing of the lungs in both species (Ridley *et al.*, 2014), whereas MUC5AC with its more complex polymers is typical for normal pig and human surface goblet cells and in all species more related to disease. The ratio of these two mucins are similar in BALF from the PPE mice and COPD suggesting that the composition of the accumulated mucus should be similar despite the large differences in normal lung of mice and men. In the mouse, more Muc5b mucin were concentrated to the mucus plugs and BALF washings of Muc5b/Muc5ac knock-out animals also suggested that Muc5b contributed most to mucus obstruction. This relative larger importance of Muc5b in mouse is also observed for normal lung clearance (Roy *et al.*, 2014). Even if we do not fully understand the details of how these two mucins contribute to mucus properties, it may suggest that the accumulated disease associated mucus has similar features in the two species.

The protein composition of the mucus accumulated in the PPE mice and COPD show large similarities to that of the inner colon mucus layer. Even more intriguing is that the structure of the accumulated mucus in COPD, CF and in the PPE-exposed mice show a striated arrangement closely resembling that of the inner mucus layer of normal colon (Johansson *et al.*, 2008). The way to observe mucus is to utilize Carnoy-fixation that better preserves the structure of the water-rich mucus. In colon, the inner mucus layer act as a barrier that physically separates the luminal bacteria from the epithelial surface (Johansson *et al.*, 2008). This separation is obtained by the MUC2 mucin forming net-like sheets that when layered on each other form a filter and thereby protect the epithelial cells. We now suggest that upon infection of the airways, the mucus which is normally easily moved by cilia, turns into an attached mucus layer resembling that in the normal colon, with similar components and structure. It is reasonable to believe that this airway mucus layer should act as in colon, i.e. as an innate defense mechanism aimed at keeping the bacteria away from the host epithelial cell surface.

Washing the PPE-exposed mice lungs with PBS partly removed the mucus plugs blocking especially the more peripheral airways. However, a similar proportion of the epithelial surface was covered by mucus without and after washing suggesting that the mucus was attached. Studies of tissue sections, also after intense washings, suggested that the mucus and the two mucins were anchored in the goblet cells. Interestingly, this is similar to what has been observed for mucus in the CF small intestine where the Muc2 mucin is not properly unfolded and thus not released from its attachment by the meprin β protease (Gustafsson *et al.* 2012; Schütte *et al.*, 2014). To test if the commonly used CF treatment, hypertonic saline, could further detach the attached mucus we treated the PPE-exposed mice two times with 7% NaCl. Hypertonic saline significantly decreased airway mucus obstruction better than PBS. However, this treatment was not better than PBS in removing the attached mucus that covered the epithelial surface. Together this makes it likely that the attachment of the mucus to the goblet cells is the main mechanism for obtaining an airway mucus layer.

The normal lung is kept relatively free of bacteria by the cilia mediated mucus escalator. However, if this fails we now suggest that a second line of defense is triggered by the formation of an attached mucus layer, similar to the normal protection of the colon epithelium. The physiological function of this airway mucus layer might thus be to protect the epithelial cells. This attached stratified mucus layer may be a normal physiological mechanism attempting to separate the bacteria from the epithelial cells to give the immune system a possibility to raise an immune response. Once the bacteria are under control, the mucus should be removed after detachment. However, this type of mucus will not be transported very efficiently by the cilia and probably has to be coughed up, a typical resolution to any lung infection. In chronic lung diseases like chronic bronchitis, COPD, CF and severe asthma, the mucus probably remains attached as shown here and although the epithelium is better protected, the bacteria retention will lead to lung deterioration. The intriguing question is now the molecular details regulating mucus attachment.

### Materials and Methods

#### Animals

Female C57BL/6N (Taconic), Muc5b^-/-^, and Muc5ac^-/-^ (Roy *et al.*, 2014) mice were used when 10 ± 2 weeks old and weighting 21 ± 3 g. Animals were housed in a specific pathogen-free facility in groups of three or four in controlled temperature (21 ± 2 °C), relative humidity (55 ± 15%) and a 12-hour light/ dark cycle. Standard chow and water were available *ad libitum*.

#### PPE administration and BALF collection

Animals under light isoflurane anesthesia were exposed to 0,312 units of porcine pancreatic elastase (PPE; Ref.45124, Sigma-Aldrich), inactivated PPE or vehicle (50 μL saline) instilled intranasally. The instillation procedure was performed at days 0 and 7. Six to eight days post final exposure, animals were euthanized with an overdose of pentobarbital sodium i.p. and the abdominal aorta was severed. To obtain BALF samples, tracheas were cannulated with a catheter (Venflon, Becton Dickinson). Lungs were manually lavaged twice with 0.8 mL of PBS for 1 min each and the 2 lavages from single animals were pooled. BALF samples and supernatants, obtained after centrifugation (500 g, 10 min, 4°C), were stored at - 80°C for later analysis. In additional experiments, the duration of each lavage was increased to 20 min and, besides PBS, hypertonic saline (7% NaCl) was tested to investigate the ability to detach airway mucus.

#### Blood cell count and inflammatory mediator analysis

Total and differential white blood cell (WBC) counts were determined using an automated cell counter (Sysmex XT-2000iV; Sysmex Corporation). Cytokine analysis in BALF supernatant was assessed using MILLIPLEX MAP Mouse Cytokine/Chemokine Magnetic Bead Panels (MCYTMAG70PMX25BK and MECY2MAG-73K; Millipore) and the Proinflammatory Panel 1 (mouse) kit (N05048A–1; Meso Scale Discovery). Epithelial growth factor (EGF) concentrations were determined by ELISA (MEG00; R&D Systems). Assays were used according to the respective manufacturer’s instructions.

#### Mucus plug isolation

For mucus plug isolation, BAL was performed with an Alcian blue solution in PBS. First, a 5% wt/vol suspension of Alcian blue 8GX (Sigma-Aldrich) in PBS was prepared. After mixing overnight (1,000 rpm) in a Thermomixer comfort (Eppendorf) at room temperature, this was centrifuged (14,000 g, 10 min, room temperature). The supernatant was further diluted 1:40 in PBS. Mucus plugs, identified as Alcian blue-stained structures in BALF, were transferred with a 10 μl tip to a dry tube, frozen in liquid N_2_ and stored at - 80°C.

#### PPE inactivation

To inactivate the serine peptidase activity of PPE, a stock solution of 50 units/ml was incubated with 1 mM PMSF overnight at 4°C. The resulting solution was diluted 1:10 in sterile saline and 1 ml was loaded in dialysis membrane tubing (Spectrum™ Spectra/Por™ 4 RC 12-14 KDa; Fisher Scientific) and dialyzed against 1 l of saline at 4 °C twice (first dialysis: 8 h; second dialysis: overnight). The inactivated PPE was concentrated using a centrifugal filter (10 kDa pore size; UFC501024; Merck Millipore). The protein concentration of the inactivated PPE was assessed by Nanodrop (A280; Thermo Scientific). The elastase activity of the inactivated PPE, active PPE (positive control) and saline were measured using an EnzChek Protease Assay Kit (Thermo Fisher Scientific). The substrate BODIPY casein was added and fluorescence, as a measure of cleavage, was read at λ_ex_477 ± 14 nm and λ_em_525 ± 30 nm every 5 min for 1 h (Fig S1). The percentage of inhibition was calculated for the last reading.

#### Histopathological analysis

After obtaining BALF, the thoracic cavity was opened. Lungs used to characterize inflammatory response and goblet cell hyperplasia were perfused through the interventricular septum with 10ml PBS and then inflated to 25 cm H_2_O static pressure by intratracheal instillation of Carnoy (60% methanol dried, 30% chloroform stabilized with ethanol and 10% glacial acetic acid 100% anhydrous). The trachea was tied and the inflated lungs removed and immersed in fixative. Lungs used to study airway mucus accumulation and plugging were directly immersed in Carnoy, without previous inflation. Lungs were embedded in paraffin and sections (3 μm), comprising both the left and the right lung lobes, were cut at the level of the tracheobronchial tree. Sections were stained with H&E to evaluate inflammation and structural alterations, with Alcian blue/periodic acid-Schiff (AB/PAS) to assess goblet cell hyperplasia and mucus accumulation or used for immunostainings.

#### Quantification of mucus accumulation

Images of mouse lung sections stained with AB/PAS were obtained at 100X magnification using a Nikon eclipse E1000 microscope and the NIS elements software (Nikon). Images of all airway sections included in an entire lung section were captured. ImageJ (National Institute of Health, Bethesda, MD) was used for image analysis. The percentage of the lumen area positively stained with AB/PAS for each airway section was measured. For each mouse, we calculated the total percentage of lumen airway obstruction (AB/PAS positive). In addition, we evaluated for each airway section the minimum Feret’s diameter (XFmin), a very robust size measure against experimental errors such as the orientation of the sectioning angle (Birrer *et al.*, 1994). The correlation of XFmin with the percentage of obstruction was assessed in PPE-exposed mice and the percentage of airway epithelial surface covered by AB/PAS-positive material was calculated.

#### Immunostainings

Lung tissue sections were dewaxed, rehydrated and microwave antigen retrieval was performed (10 mM citrate buffer pH 6). Samples were blocked for 20 min at room temperature with 5% inactivated goat or donkey serum, according to the species in which the secondary antibodies were produced. Thereafter, sections were incubated overnight at 4 °C with a mouse anti-Muc5ac antibody (clone 45M1; M5293, Sigma-Aldrich; 1:1,000), a rabbit anti-Muc5b antibody (Mehmet Kesimer, Chapell Hill, NCU; 1:2,000), a rabbit anti-Clca1 antibody (ab46512, Abcam; 1:1,000), a rabbit anti-Fcgbp antibody (HPA003517, Atlas Antibodies; 1:100), or a mouse monoclonal anti-tubulin antibody (T6793, Sigma-Aldrich; 1:4,000). In case of single stains, slides were then incubated for 2 h at room temperature with goat anti-mouse IgG or donkey anti-rabbit IgG secondary antibodies conjugated to Alexa Fluor 488 or 555 (Life Technologies; 1:1,000). For co-localization studies, slides were washed, blocked, incubated with a second primary antibody, using the same conditions described above, and then incubated for 2 h at room temperature with the corresponding fluorescently conjugated antibodies (1:1,000). Nuclei were counterstained for 10 min using 1 μg/ml Hoechst 34580 (H21486, ThermoFisher Scientific) and mounted with Prolong Gold Antifade mounting medium (P36930, Life Technologies). The specificity of the primary antibodies used for Muc5ac and Muc5b detection has been previously confirmed by others (Lidell *et al.*, 2008;Zhu *et al.*, 2008). Concerning the antibodies for Clca1 and Fcgbp, we confirmed that none of them stained sections of colon or PPE-exposed lungs in corresponding knock out mice. Non-specific binding of the secondary antibody was ruled out by omitting the primary antibody. Whole lung sections were scanned using an Axio Scan.Z1 scanner with a 20X Plan-Apochromat 0.8 NA dry objective and ZEN 2.1 software (Carl Zeiss). High magnification images were obtained using a LSM700 Axio Examiner Z1 confocal microscope (Carl Zeiss) with the ZEN 2010 software (Carl Zeiss). Fluorescence intensity per μm^2^ was measured with ImageJ and expressed as the relative change compared to the median intensity in saline-treated controls. Analysis of all data was performed in a blinded manner.

#### Transmission electron microscopy

Mouse lungs were fixed in modified Karnovsky’s fixative (2% paraformaldehyde, 2.5% glutaraldehyde in 0.05 M, sodium cacodylate buffer, pH 7.2) for 24 h. One lung lobe was divided into 9 transverse sections and every other section starting from the second section was processed further for electron microscopy. Samples were sequentially stained using 1% OsO_4_ for 4 h, 1% tannic acid for 3 h and 1% uranyl acetate overnight then dehydrated and embedded in epoxy resin (Agar 100, Agar Scientific). Electron microscopy was conducted on 50 nm sections cut using an Ultracut E (Reichert) microtome and collected on mesh copper support grids. The sections were contrasted using lead citrate and tannic acid and images were acquired using a LEO 912 transmission electron microscope (Carl Zeiss) operated at an accelerating voltage of 120 kV.

#### Isolation of Epithelial Cells

After obtaining BALF, lungs were washed twice through the intratracheal cannula with 0.8 ml PBS and the liquid discarded. Thereafter, epithelial cells were isolated using a predisgestion buffer (20 mM EDTA and 1 mM DTT in HBSS without Ca^2+^ or Mg^2+^ supplemented with 10 mM HEPES). Initially, 0.8 ml of pre-warmed predisgestion buffer was instilled through the intratracheal cannula. Lungs were immersed in pre-warmed PBS and placed in a water bath at 37°C, where they remained during the whole procedure. Lungs were then washed with 0.8 ml of pre-warmed predigestion buffer once at 15 and 30 min and 4 times at 45 min and 1 h after the first instillation, respectively. After each lavage, the buffer aspirated from the lungs containing epithelial cells in suspension was transferred to a tube kept on ice. All the lavages obtained from 3 animals belonging to the same group were pooled to obtain each epithelial cell sample. The cell suspension was vortexed for 2 min and centrifuged (500 g, 10 min, 4°C). The supernatant was discarded and the cell pellet resuspended in 2 ml HBSS and kept on ice. Erythrocytes were lysed using the ACK Lysing Buffer (Gibco, Thermo Fisher Scientific). Afterwards, cell pellets were incubated in 1 ml of collagenase type I (2mg/ml; 17100-017, Invitrogen) and DNAse I (40 U/ml; D-4513, Sigma-Aldrich) in DMEM/F12 with HEPES for 30 min at 37°C. After enzymatic digestion, cells were washed with 10 ml HBSS, centrifuged (500 g, 10 min, 4°C) and resuspended in 2 ml HBSS. The cell suspension was passed through a 30 µm mesh (Miltenyi Biotec) to obtain a single-cell suspension. A sequential negative selection of epithelial cells was performed using MACS separation reagents and instruments (Miltenyi Biotec Inc). Samples were first depleted of CD45 positive cells and then of Ly6G positive cells, using two different MACS LD Depletion column, microbeads and MidiMACS™ magnets (all from Miltenyi Biotec). The procedure was performed according to manufacturer’s instructions with the exception that columns were washed 3 times instead of 2, to maximize the recovery of unlabeled cells, and that the incubation with Anti-Ly-6G-Biotin beads was increased from 10 to 30 min, to improve neutrophil depletion. At the end of the procedure, the magnetically unlabeled cell fraction containing epithelial cells was collected. Ice-cold PBS was added to a total volume of 14 ml, samples were centrifuged (500 g, 10 min, 4°C) and the supernatant discarded. Cells were resuspended in 1 ml PBS, transferred to MCT–175–L–C tubes (Axygen) and centrifuged (1000 g, 5 min, 4°C). After the supernatant was completely removed, pellets were frozen in liquid N_2_ and stored at – 80°C.

#### Mass spectrometry

Mice BALF (50 μl) and mucus plug samples were incubated overnight at 37°C in reduction buffer (6 M guanidinium hydrochloride (GuHCl), 0.1 M Tris/ HCl pH 8.5 (Merck), 5 mM EDTA, M DTT, Merck) followed by a modified filter-aided sample preparation (FASP) protocol using 6 M GuHCl (van der Post and Hansson, 2014). Briefly, proteins were alkylated on 10 kDa cut-off filters and subsequently digested for 4 h with LysC (Wako Chemicals) followed by an overnight trypsin (Promega) digestion. Heavy peptides (JPT Peptide Technologies) for Muc5b and Muc5ac absolute quantification (5 peptides for each protein, 100 fmol each; Table S7) were added before trypsin digestion. Epithelial cells for proteomics analysis were prepared according to the FASP protocol (Wisniewski *et al.*, 2009). Briefly, cells were lysed by boiling in 4% SDS, buffer was exchanged to 8 M urea on 30 kDa cut-off filters and proteins were alkylated and digested for 4 h with endopeptidase LysC (Wako Chemicals), followed by overnight digestion with trypsin (Promega). Human BALF (180 µl) was processed as mice BALF except heavy labeled peptides (JPT Peptide Technologies) for 15 human proteins were added (100 fmol each; Table S7).

Peptides released from the filter by centrifugation were cleaned with StageTip C18 columns as previously described (Rappsilber *et al.*, 2007). Nano-Liquid-Chromatography Mass-spectrometry (NanoLC–MS/MS) was performed on an EASY-nLC system (Thermo Scientific), connected to a Q-Exactive Hybrid Quadrupole-Orbitrap Mass Spectrometer (Thermo Scientific) through a nanoelectrospray ion source. Peptides from mucus plugs and BALF samples were separated with in-house packed C18 reverse-phase column (150 × 0.075 mm inner diameter, C18-AQ 3 μm) by 60-minute gradient from 5 to 40% B (A: 0.1% formic acid, B: 0.1% formic acid/80% acetonitrile 200 nl/min). Peptides from epithelial cell samples were separated with heated (50°C) C18 reverse-phase EASY-Spray column (500 × 0.050 mm, PepMap RSLC, 2 μm, 100 Å) by a 120-minute gradient from 2-25% B, followed by 10-minutes increase to 40% B. Full mass spectra were acquired from m/z320–1,600, for BALF and plugs, or 400–1,600 m/z, for epithelial cell pools, with resolution of 70,000. Up to twelve most intense peaks (charge state ≥ 2) were fragmented and tandem mass spectrum was acquired with a resolution of 35,000 and automatic dynamic exclusion for BALF and mucus plugs samples and with a resolution of 17,500 and dynamic exclusion 10 s for epithelial cells. Human BAL fluid samples were analyzed by acquiring full spectrum from m/z 350–1, 600 m/z with resolution of 70,000, up to ten most intense peaks were fragmented and tandem mass spectrum was acquired with a resolution of 17,500 and automatic dynamic exclusion of 10 s. For absolute quantification a separate targeted mass spectrometry method was used where only precursors and their fragments of the heavy and corresponding light peptides were scanned with a resolution of 35,000.

#### Proteomics data analysis

Proteomics data were analyzed with the MaxQuant program (version 1.4.1) (Cox and Mann, 2008), by searching against the mouse UniProt protein database (downloaded July 23, 2015) supplemented with an in-house database containing all the mouse mucin sequences (http://www.medkem.gu.se/mucinbiology/databases/) or human Uniprot protein database(downloaded July 23, 2015) supplemented with an in-house database containing all the human mucin sequences(http://www.medkem.gu.se/mucinbiology/databases/). Searches were performed with full tryptic specificity, maximum 2 missed cleavages, precursor tolerance of 20 ppm in the first search used for recalibration, followed by 7 ppm for the main search and 0.5 Da for fragment ions. Carbamidomethylation of cysteine was set as a fixed modification and methionine oxidation and protein N-terminal acetylation were set as variable modification. The required false discovery rate (FDR) was set to 1% both at the peptide and protein levels and the minimum required peptide length was set to six amino acids. Proteins were quantified based on MaxQuant label-free quantification (LFQ) option using minimum two peptides for quantification. Data filtering, clustering and principal clustering analysis (PCA) was performed with Perseus software (version 1.5.6.0). The data was filtered based on the presence of a protein group in at least three out of six biological replicated for BAL fluid, four out of seven replicates for epithelial cells, six out of 12 replicates for plugs. Missing values were replaced from normal distribution with default values. In the final qualitative lists albumin was excluded as contaminant (otherwise contributing up to 70% of total intensities). Clustering and heatmaps were performed using Euclidean distance, average linkage, preprocessing with k-means for rows and columns. For principal component analysis, default parameters were used, number of components 5, cutoff method Benjamin-Hochberg with FDR 0.05. The mass spectrometry proteomics data were deposited to the Proteome Xchange Consortium (http://proteomecentral.proteomexchange.org/) via the PRIDE partner repository (https://www.ebi.ac.uk/pride/archive/) with the dataset identifier PXDXXXXX. Absolute quantification proteins that had heavy labeled standard peptides (Table S7) was performed with the Skyline program (version 3.6.0.1) (MacLean *et al.*, 2010). Peptides were quantified using isotopically labelled analogues and median peptide concentration was reported as protein concentration.

#### Patient samples

BALF samples were collected from healthy non-smokers, healthy smokers, and patients with diagnosed COPD. The patients included are gathered in Table S5

#### Study approval

Care and use of animals was undertaken in compliance with the European Community Directive 2010/63/EU. Animal experiments were approved by the local ethics committee in Gothenburg with the ethical permit numbers 100-2014 and 75–2015. Human samples were acquired with approval by the local human research ethics committee with the permit number 543–11. All human subjects gave written informed consent prior to inclusion and all investigations were performed in accordance with the Declaration of Helsinki.

#### Statistics

Results are presented as median ± interquartile range, mean ± SEM. For comparing two independent groups, we used the Mann-Whitney U test and for comparison of multiple groups we used Kruskal-Wallis and Dunn’s post-test to correct for multiple comparisons. The number of biological replicates is indicated by n number of animals or samples when several animals were pooled to obtain one data point. Significance was defined as P < 0.05.

## Acknowledgments

The work was funded by AstraZeneca, the European Research Council ERC (694181), National Institute of Allergy and Infectious Diseases (2U01AI095473, the content is solely the responsibility of the authors and does not necessarily represent the official views of the NIH), The Knut and Alice Wallenberg Foundation, The Swedish Research Council, The Swedish Cancer Foundation, IngaBritt and Arne Lundberg Foundation, Sahlgrenska University Hospital (ALF), Wilhelm and Martina Lundgren’s Foundation, The Cystic Fibrosis Foundation (CFF), Swedish CF Foundation, Erica Lederhausen’s Foundation, Lederhausen’s Center for CF Research at Univ. Gothenburg, Magnus Bergvall´s foundation and the Swedish Heart-Lung Foundation. We thank Dr. Mehmed Kesimer (University of North Carolina, Chapell Hill, NC, USA) for the anti-Muc5b antibody. We also acknowledge the Centre for Cellular Imaging at the University of Gothenburg, Lisa Jinton for cytokine analyses, Sofia Lundin for handling of human tissues, and Botilda Lindberg and Susanne Arlbrandt for animal work.

## Author contributions

JAF–B, A;E, AÅ and GCH designed the study. JAF–B, LA, AE, AMR–P, BM–A, ES, and SJ conducted the experiments. SJ, JR, CM, CE, and AÅ provided compounds and materials. DS provided patient samples. JAF–B, AE and GCH wrote the manuscript. All authors approved the manuscript.

## For more information

Supplementary information accompanying this paper at doi:

## References

1 Bateman JR, Pavia D, Sheahan NF, Agnew JE, and Clarke SW (1983) Impaired tracheobronchial clearance in patients with mild stable asthma. Thorax, 38, 463–467.

2 Bergström JH, Birchenough GMH, Katona G, Schroeder BO, Schutte A, Ermund A, Johansson MEV, and Hansson GC (2016) Gram-positive bacteria are held at a distance in the colon mucus by the lectin-like protein ZG16. Proc Natl Acad Sci USA, 113, 13833–13838.

3 Bingle L, Wilson, Musa M, Araujo B, Rassl D, Wallace WA, LeClair EE, Mauad T, Zhou Z, Mall MA, and Bingle CD (2012) BPIFB1 (LPLUNC1) is upregulated in cystic fibrosis lung disease. Histochem Cell Biol, 138, 749–758.

4 Birrer P, McElvaney NG, Rudeberg A, Sommer CW, Liechti-Gallati S, Kraemer R, Hubbard R, and Crystal RG (1994) Protease-antiprotease imbalance in the lungs of children with cystic fibrosis. Am J Respir Crit Care Med, 150>, 207–213.

5 Conlon TM, Bartel J, Ballweg K, Gunter S, Prehn C, Krumsiek J, Meiners S, Theis FJ, Adamski J, Eickelberg O, and Yildirim AO (2016) Metabolomics screening identifies reduced L-carnitine to be associated with progressive emphysema. Clin Sci (Lond), 130, 273–287.

6 Cox J and Mann M(2008) MaxQuant enables high peptide identification rates, individualized p.p.b.-range mass accuracies and proteome-wide protein quantification. Nature Biotechnol, 26, 1367–1372.

7 Ermund A, Meiss LN, Rodriguez-Pineiro AM, Bähr A, Nilsson HE, Trillo-Muyo S, Ridley C, Thornton DJ, Wine JJ, Hebert H, Klymiuk N, and Hansson GC (2017) The normal trachea is cleaned by MUC5B mucin bundles from the submucosal glands coated with the MUC5AC mucin. Biochem Biophys Res Commun, 492, 331–337.

8 Fu XW, Song PF, and Spindel ER (2015)) Role of Lynx1 and related Ly6 proteins as modulators of cholinergic signaling in normal and neoplastic bronchial epithelium. Int Immunopharmacol, 29, 93–98.

9 Gao J, Ohlmeier S, Nieminen P, Toljamo T, Tiitinen S, Kanerva T, Bingle L, Araujo B, Ronty M, Hoyhtya M, Bingle CD, Mazur W, and Pulkkinen V(2015) Elevated sputum BPIFB1 levels in smokers with chronic obstructive pulmonary disease: a longitudinal study. Am J Physiol Lung Cell Mol Physiol, 309, L17–L26.

10 Gross P, Babyak MA, Tolker E, and Kaschak M (1964) Enzymatically produced pulmonary emphysema; a preliminary report. J Occup Med, 6, 481–484.

11 Gustafsson JK, Ermund A, Ambort D, Johansson MEV, Nilsson HE, Thorell K, Hebert H, Sjovall H, and Hansson GC (2012) Bicarbonate and functional CFTR channel is required for proper mucin secretion and link Cystic Fibrosis with its mucus phenotype. J Exp Med, 209, 1263–1272.

12 Hoenderdos K and Condliffe A (2013) The neutrophil in chronic obstructive pulmonary disease. Am J Respir Cell Mol Biol, 48, 531–539.

13 Hogg JC, Chu F, Utokaparch S, Woods R, Elliott WM, Buzatu L, Cherniack RM, Rogers RM, Sciurba FC, Coxson HO, and Pare PD (2004) The Nature of Small-Airway Obstruction in Chronic Obstructive Pulmonary Disease. N Engl J Med, 350, 2645–2653.

14 Johansson MEV, Phillipson M, Petersson J, Holm L, Velcich A, and Hansson GC (2008) The inner of the two Muc2 mucin dependent mucus layers in colon is devoid of bacteria. Proc Natl Acad Sci USA, 105, 15064–15069.

15 Johansson MEV, Thomsson KA, and Hansson GC (2009) Proteomic Analyses of the Two Mucus Layers of the Colon Barrier Reveal That Their Main Component, the Muc2 Mucin, Is Strongly Bound to the Fcgbp Protein. J Proteome Res, 8, 3549–3557.

16 Komi DE, Kazemi T, and Bussink AP (2016) New Insights Into the Relationship Between Chitinase-3-Like-1 and Asthma. Curr Allergy Asthma Rep, 16, 57.

17 Kung TT, Jones H, Adams GK, III, Umland SP, Kreutner W, Egan RW, Chapman RW, and Watnick AS (1994) Characterization of a murine model of allergic pulmonary inflammation. Int Arch Allergy Immunol, 105, 83–90.

18 Kuperman DA, Huang X, Koth LL, Chang GH, Dolganov GM, Zhu Z, Elias JA, Sheppard D, and Erle DJ (2002) Direct effects of interleukin-13 on epithelial cells cause airway hyperreactivity and mucus overproduction in asthma. Nat Med, 8, 885–889.

19 Lee R, Han K, and Han J (2015) rbCLCA1 is a putative metalloprotease family member: localization and catalytic domain identification. Amino Acids, 1–14.

20 Lidell ME, Bara J, and Hansson GC (2008) Mapping of the 45M1 epitope to the C-terminal cysteine-rich part of the human MUC5AC mucin. FEBS J, 275, 481–489.

21 Lukacs NW (2001)Role of chemokines in the pathogenesis of asthma. Nat Rev Immunol, 1, 108–116.

22 MacLean B, Tomazela DM, Shulman N, Chambers M, Finney GL, Frewen B, Kern R, Tabb DL, Liebler DC, and MacCoss MJ (2010) Skyline: an open source document editor for creating and analyzing targeted proteomics experiments. Bioinformatics, 26, 966–968.

23 Madsen J, Mollenhauer JF, and Holmskov U (2010) Review: Gp-340/DMBT1 in mucosal innate immunity. Innate Immun, 16, 160–167.

24 Matsui H, Grubb BR, Tarran R, Randell SH, Gatzy JT, Davis CW, and Boucher RC (1998) Evidence for periciliary liquid layer depletion, not abnormal ion composition, in the pathogenesis of cystic fibrosis airways disease. Cell, 95, 1005–1015.

25 Ohlmeier S, Nieminen P, Gao J, Kanerva T, R+Ânty M, Toljamo T, Bergmann U, Mazur W, and Pulkkinen V (2016) Lung tissue proteomics identifies elevated transglutaminase 2 levels in stable chronic obstructive pulmonary disease. American Journal of Physiology - Lung Cellular and Molecular Physiology, 310, L1155.

26 Podolsky DK, Lynchdevaney K, Stow JL, Oates P, Murgue B, Debeaumont M, Sands BE, and Mahida YR (1993) Identification of Human Intestinal Trefoil Factor - Goblet Cell-Specific Expression of a Peptide Targeted for Apical Secretion. (vol 268, pg 6694, 1993) J Biol Chem, 268, 12230.

27 Rappsilber J, Mann M, and Ishihama Y (2007) Protocol for micro-purification, enrichment, pre-fractionation and storage of peptides for proteomics using StageTips. Nat Protoc, 2, 1896–1906.

28 Recktenwald CV and Hansson GC (2016) The reduction-insensitive bonds of the MUC2 mucin are isopeptide bonds. J Biol Chem, 291, 13580–13590.

29 Regnis JA, Robinson M, Bailey DL, Cook P, Hooper P, Chan HK, Gonda I, Bautovich G, and Bye PT (1994) Mucociliary clearance in patients with cystic fibrosis and in normal subjects. Am J Respir Crit Care Med, 150, 66–71.

30 Ridley C, Kouvatsos N, Raynal BD, Howard M, Collins RF, Desseyn JL, Jowitt TA, Baldock C, Davis CW, Hardingham TE, and Thornton DJ (2014) Assembly of the Respiratory Mucin MUC5B: a new model for gel-forming mucin. J Biol Chem, 289, 16409–16420.

31 Rodriguez-Pineiro AM, Bergstrom JH, Ermund A, Gustafsson JK, Schutte A, Johansson MEV, and Hansson GC (2013) Studies of mucus in mouse stomach, small intestine, and colon. II. Gastrointestinal mucus proteome reveals Muc2 and Muc5ac accompanied by a set of core proteins. Am J Physiol Gastroint Liver Physiol, 305, G348–G356.

32 Roy MG, Livraghi-Butrico A, Fletcher AA, McElwee MM, Evans SE, Boerner RM, Alexander SN, Bellinghausen LK, Song AS, Petrova YM, Tuvim MJ, Adachi R, Romo I, Bordt AS, Bowden MG, Sisson JH, Woodruff PG, Thornton DJ, Rousseau K, De la Garza MM, Moghaddam SJ, Karmouty-Quintana H, Blackburn MR, Drouin SM, Davis CW, Terrell KA, Grubb BR, ’Neal WK, Flores SC, Cota-Gomez A, Lozupone CA, Donnelly JM, Watson AM, Hennessy CE, Keith RC, Yang IV, Barthel L, Henson PM, Janssen WJ, Schwartz DA, Boucher RC, Dickey BF, and Evans CM (2014) Muc5b is required for airway defence. Nature, 505, 412–416.

33 Schütte A, Ermund A, Becker-Pauly C, Johansson MEV Rodriguez-Pineiro AM, Bäckhed F, Müller S, Lottaz D, Bond JS, and Hansson GC (2014) Microbial Induced Meprin beta Cleavage in MUC2 Mucin and Functional CFTR Channel are Required to Release Anchored Small Intestinal Mucus. Proc Natl Acad Sci USA, 111), 12396–12401.

34 Shapiro SD (2000) Animal models for COPD. Chest, 117, 223S–227S.

35 Smaldone GC, Foster WM, O’Riordan TG, Messina MS, Perry RJ, and Langenback EG (1993) Regional impairment of mucociliary clearance in chronic obstructive pulmonary disease. Chest, 103, 1390–1396.

36 Thornton DJ, Rousseau K, and McGuckin MA (2008) Structure and Function of the Polymeric Mucins in Airways Mucus. Ann Rev Physiol, 70, 459–486.

37 van der Post S and Hansson GC (2014) Membrane protein profiling of human colon reveals distinct regional differences. Mol Cell Proteomics, 13, 2277–2287.

38 Vestbo J, Hurd SS, Agusti AG, Jones PW, Vogelmeier C, Anzueto A, Barnes PJ, Fabbri LM, Martinez FJ, Nishimura M, Stockley RA, Sin DD, and Rodriguez-Roisin R (2013) Global strategy for the diagnosis, management, and prevention of chronic obstructive pulmonary disease: GOLD executive summary. Am J Respir Crit Care Med, 187, 347–365.

40 Wiede A, Jagla W, Welte T, Kohnlein T, Busk H, and Hoffmann W (1999) Localization of TFF3, a new mucus-associated peptide of the human respiratory tract. Am J Respir Crit Care Med, 159, 1330–1335.

41 Wisniewski JR, Zougman A, Nagaraj N, and Mann M (2009) Universal sample preparation method for proteome analysis. Nat Methods, 6, 359–362.

41 Yamamoto C, Yoneda T, Yoshikawa M, Fu A, Tokuyama T, Tsukaguchi K, and Narita N (1997) Airway inflammation in COPD assessed by sputum levels of interleukin-8. Chest, (112), 505-510.

42 Zhu Y, Ehre C, Abdullah LH, Sheehan JK, Roy M, Evans CM, Dickey BF, and Davis CW(2008) Munc13-2-/- baseline secretion defect reveals source of oligomeric mucins in mouse airways. J Physiol, 586, 1977–1992.

